# Hecatomb: An End-to-End Research Platform for Viral Metagenomics

**DOI:** 10.1101/2022.05.15.492003

**Authors:** Michael J. Roach, Sarah J. Beecroft, Kathie A. Mihindukulasuriya, Leran Wang, Anne Paredes, Kara Henry-Cocks, Lais Farias Oliveira Lima, Elizabeth A. Dinsdale, Robert A. Edwards, Scott A. Handley

## Abstract

**Background:** Analysis of viral diversity using modern sequencing technologies offers extraordinary opportunities for discovery. However, these analyses present a number of bioinformatic challenges due to viral genetic diversity and virome complexity. Due to the lack of conserved marker sequences, metagenomic detection of viral sequences requires a non-targeted, random (shotgun) approach. Annotation and enumeration of viral sequences relies on rigorous quality control and effective search strategies against appropriate reference databases. Virome analysis also benefits from the analysis of both individual metagenomic sequences as well as assembled contigs. Combined, virome analysis results in large amounts of data requiring sophisticated visualization and statistical tools.

**Results:** Here we introduce Hecatomb, a bioinformatics platform enabling both read and contig based analysis. Hecatomb integrates query information from both amino acid and nucleotide reference sequence databases. Hecatomb integrates data collected throughout the workflow enabling analyst driven virome analysis and discovery. Hecatomb is available on GitHub at https://github.com/shandley/hecatomb.

**Conclusions:** Hecatomb provides a single, modular software solution to the complex tasks required of many virome analysis. We demonstrate the value of the approach by applying Hecatomb to both a host-associated (enteric) and an environmental (marine) virome data set. Hecatomb provided data to determine true- or false-positive viral sequences in both data sets and revealed complex virome structure at distinct marine reef sites.

## Background

Viruses parasitize host cell molecular processes and as a result alter host (prokaryotic and eukaryotic) cell physiology. Virus-host interactions can influence organismal physiology and environmental ecosystems. Viruses are also the most dominant entity on the planet with current global estimates as high as 10^31^ viral particles [1,2], and they are omnipresence in all cellular life forms [3]. As such they exert significant influence on their surroundings.

The effect of viruses on human life and society are dramatically demonstrated through phenomena such as global pandemics. However, the true burden of viruses on human health is incredibly varied in terms of breadth and severity. There are many well-known acute viral diseases such as the “common cold” (rhinoviruses, adenoviruses and enteroviruses) [4] which cause tremendous amounts of morbidity, but limited mortality. In contrast, chronic Epstein-Barr virus (EBV) infection has recently been associated with the onset of multiple sclerosis [5]. Consequential virus-host interactions are not limited to humans. For example, Geminivirus infection of plants has resulted in nearly US$2 billion loss in African cassava production [6]. Similar foot-and-mouth disease virus (FMDV), a highly contagious disease of cloven-hoofed animals, is widespread in Africa with an annual US$2.3 billion negative impact on livestock [7]. Virus ‘spillover’ infection from animal to human (“zoonotic” viruses) is unfortunately an all too regular event [8]. Viral zoonosis from viruses such as SARS-CoV-2, monkeypox, Ebola and Zika viruses have tremendous negative impacts on human health and society and new zoonotic viruses are constantly emerging presenting a persistent threat to human health [9].

Viral assemblages, often referred to as *viromes*, are also associated with human health and disease [10]. Stool samples from patients with inflammatory bowel disease (IBD) suffer dysbiosis of microbial populations, having expanded numbers of bacteriophage (hereafter *phage*) from the order Caudovirales [11–15]. Enteric vertebrate virus expansion occurs in both rhesus macaques and humans with acquired immunodeficiency syndrome (AIDS) [16,17]. Thus, health is not only influenced by infection with single viruses, but also viromes. A comprehensive virus analysis workflow enables the analysis of both.

Viruses also influence global ecosystems. For example, the release of intracellular iron and sulphur from bacteria following lytic phage infection releases nutrients used by phytoplankton into marine environments via a mechanism called a *viral shunt* [18]. These phytoplankton are in-turn eaten by higher trophic levels altering the entire marine food web. Many other aquatic nutrient cycling and biogeochemical processes are attributed, both directly and indirectly, to viral modification of prokaryotic and protistan assemblages [19–23]. Terrestrial environment carbon and nutrient cycling are also influenced by bacteriophage [24–26]. Viral modification of both aquatic and terrestrial ecosystems underlies the importance of environmental virome studies to comprehensively understand climate, ecology and production. Virome analysis tools should enable detailed interrogation of viruses from any ecosystem broadening our understanding of the global virome.

Metagenomic sequencing offers a powerful tool to study viral diversity [27]. However, there are currently many challenges associated with viral metagenomics. While viruses are the most abundant and diverse biological entity on the planet, they represent a minority of reference genomes in GenBank, largely due to difficulties associated with studying them [28]. Recent efforts to populate new viral genomes into reference databases are slowly closing this gap, and have yielded 10s to 100s of thousands of novel metagenome-assembled viral genomes [29–35]. Other efforts have yielded many new high-quality viral genomes by combining the laborious and time-consuming experimental work with student-learning outcomes [36]. Despite these efforts, there is still a vast amount of sequence information that remains taxonomically or functionally ill-defined. These sequences are regularly referred to as “viral dark matter” and poses a significant barrier to the annotation of viral sequences from metagenomic data (reviewed in [37]). The success of viral annotation is directly impacted by the size and diversity of the reference database. Sensitive search algorithms are better able to identify viral sequences that are only distantly related to reference database sequences. More diverse databases improve viral sequence annotation, but larger databases are less conducive to these high sensitivity searches due to increased computational requirements. Database limitations are further amplified when deciding to query sequences against amino acid or nucleotide reference databases. Translated searches to amino acid databases offer superior sensitivity, however, limiting searches solely to amino acid databases risks missing sequences only available in nucleotide databases.

Another challenge to interpretation of reference based sequence annotation is that viral metagenomes are often plagued with false positive classifications [38–40]. Viruses share sequence homology with other domains of life, including ‘stolen’ genes incorporated from their hosts’ genomes, and repetitive or low-complexity regions that are also found in other organisms, such as insertion elements or transposons [38–40]. These sequences are present in reference databases and can result in false-classifications due to shared sequence similarity across taxonomies. The presence of false-positive classifications may influence data interpretation. For instance, mis-classification of viral sequences in clinical samples could lead to incorrect hypotheses about virus-disease associations or patient diagnosis. Similarly, an increased false-positive rate in any environment could lead to over-estimates of species richness and diversity. False-positive taxonomic assignments are largely unavoidable without highly-curated databases which require tremendous resources and time at the risk of missing newly discovered viruses which have yet to make there way through the curation process. Thus, it is important for virome analysis bioinformatic tools to provide a system to classify the quality of taxonomic assignments empowering researchers to make informed decisions.

Here we present Hecatomb, a bioinformatics platform designed to address many of these issues. Hecatomb performs rigorous quality control followed by tiered taxonomic assignment using MMseqs2 querying sequences against virus-specific and trans-kingdom amino acid and nucleotide reference databases [41]. Hecatomb also performs metagenomic assembly and contig taxonomic classification providing simultaneous analysis of both read and contig based viral annotations. While hecatomb provides pre-compiled databases and recommended settings, it is easily customizable and extensible. The primary output of Hecatomb is a comprehensive annotation table containing data generated throughout the workflow that is designed to be easily merged with sample data for visualisation and statistical analysis. Hecatomb has been successfully applied to several viral metagenomics projects and has accelerated the discovery of novel viruses and characterisation of viral populations [42–46].

Hecatomb is open-source with the project files hosted on GitHub at github.com/shandley/hecatomb [47], with full support available using GitHub issues. Documentation and training vignettes are available at hecatomb.readthedocs.io. Documentation covers installing and optional configuration of the software; detailed information including databases, and output files; advanced usage cases; an FAQ; and a tutorial covering some example analyses of the results. Hecatomb is available for installation from the Bioconda [48] and is distributed under a permissive MIT licence. Bioconda package information for Hecatomb is available at anaconda.org/bioconda/hecatomb.

## Implementation

An overview of the Hecatomb pipeline is shown in Figure 1. Hecatomb processes reads through four key modules (Figure 1A). First (module 1), reads are preprocessed to remove low-quality or contaminating sequences (low-quality sequence, primers, adapters, host, common laboratory contaminants and duplicates). Second, preprocessed reads are passed through both a read-based analysis and an assembly module (modules 2 and 3). For taxonomic assignment, Hecatomb uses preprocessed databases (Supplementary Methods). The final module (module 4) combines information obtained from both the read-based and assembly modules. Results are stored throughout each module, primarily as tab-separated value (tsv) files for universal compatibility and easy data analysis with any framework (e.g. Python, R, Bash, Excel). Emphasis is placed on data preservation at each stage to provide analysts with as much detail as possible to inform interpretation of results.

**Figure 1:**
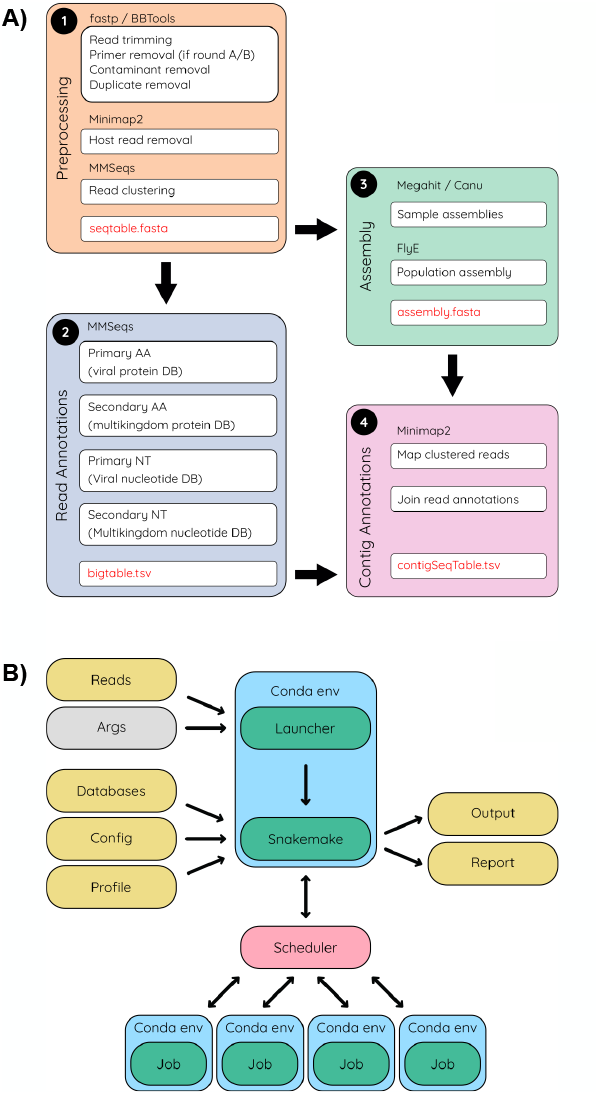
Hecatomb pipeline and implementation. **(A)** The Hecatomb pipeline is divided into four modules. Sequencing reads for each sample undergo preprocessing and clustering (*orange*); quality trimmed reads for each sample undergo assembly and assemblies for each sample are coalesced into a single assembly (*green*); clustered reads undergo annotation using viral and multi-kingdom protein databases and clustered reads not annotated by the protein search are annotated using viral and multi-kingdom nucleotide databases (*blue*); read-based annotations are combined with the assembly to provide contig annotations (*pink*). The assembly stages– *green* and *pink*–can optionally be skipped. **(B)** Hecatomb takes in command line arguments, data, configuration parameters and outputs both results for analysis and run information. Hecatomb interacts with the job scheduler in high-performance computing (HPC) environments. Hecatomb distributes individual tasks to the job queue. Command-line arguments, *grey*; files, *yellow*; Conda environments, *blue*; scripts/programs, *green*; workload manager, *pink*.

Hecatomb is installed via Conda and it makes liberal use of Conda environments to ensure portability and ease of installation (Fig 1B). All required and optional software dependencies are summarised in Table S1 [41,49–59]. Users need only install Conda which Hecatomb uses to automatically install all dependencies. Conda environments for jobs are created automatically by Snakemake [60] [49]. The use of isolated Conda environments for Hecatomb and the individual pipeline jobs minimises package version conflicts, minimises overhead when rebuilding environments for updated dependencies, and allows maintenance and customisation of different versions of Hecatomb and its dependencies without interacting with installed programs and system modules.

A custom built launcher script is included to make running the pipeline as simple as possible. The launcher populates the required file paths, the default configuration, and offers a convenient way to modify parameters and customise options. The Snakemake command generated and runtime configuration is printed to the terminal window for the user’s reference. Accessory scripts are also available from this launcher for installing reference databases, as well as adding custom host genomes, and combining results from multiple analyses.

Hecatomb is able to be deployed on an high-performance computing (HPC) cluster and has makes use of Snakemake profiles for cluster job schedulers (e.g. Slurm, SGE, etc.). Snakemake uses profiles to submit pipeline jobs to the job scheduler and monitor their progress. Although optional, using the scheduler is highly recommended as it allows for more efficient use of HPC resources compared to submitting the whole Hecatomb pipeline as a local job. Profiles can be created manually, but Hecatomb has been designed for compatibility with the official Cookiecutter (https://github.com/cookiecutter/cookiecutter) profiles for Snakemake (https://github.com/Snakemake-Profiles/doc).

### Sequence data preprocessing

Hecatomb can process both single and paired-end Illumina or MGI sequencing reads as well as long-read technology from PacBio and Oxford Nanopore platforms. Hecatomb can also process sequences obtained from other library types with minor modifications to the Hecatomb configuration file and by supplying library specific adapters or primer sequences. A preprocessing module is also available for sequencing utilising the round A/B library protocol for viral metagenomics [61]. The round A/B library protocol enables sequencing of all types of viral genomes (single and double stranded RNA and DNA viruses) and requires the use of a combination of phased PCR primers. The preprocessing module in Hecatomb removes these non-biological sequence contaminants, along with additional common laboratory sequence contaminants in the UniVec database [62].

For host-associated samples (e.g. stool, saliva, skin swabs from humans or mucus from corals) Hecatomb implements a host-sequence removal strategy using Minimap2 and a host reference genome specifically optimised to avoid removing potential viral sequences [56]. To remove all potentially viral sequences in reference genomes all viral genomes from the National Center for Biotechnology Information (NCBI) viral assembly database (ncbi.nlm.nih.gov/assembly/?term=viruses) were downloaded and computationally split into short fragments with an average length of 85 bases sharing a 30 base overlap using shred.sh from the BBTools suite [52]. Shredded viral sequences were mapped and masked from host-reference genomes using bbmap.sh requiring a minimum identity of 90% and at most, 2 insertions and deletions. In addition, low-entropy sequences were masked from host genomes (entropy = 0.5) using bbmask.sh. This process results in a set of host-associated reference genomes masked of ‘virus-like’ and low-entropy sequences, limiting the likelihood that a real viral sequence will be removed. Pre-computed masked reference genomes for the following host genomes: human, mouse, rat, camel, *Caenorhabditis elegans*, dog, cow, macaque, mosquito, pig, rat and tick are available within Hecatomb using the --host flag. Scripts are provided to generate new masked host genomes.

For the final stage of the preprocessing module, Hecatomb removes sequence redundancy by clustering each sample using linclust [63]. Clustering sequences reduces the number of sequences requiring taxonomic classification to a single, representative sequence from a cluster of similar sequences. Sequences are clustered requiring a minimum sequence identity of 97% and 80% alignment coverage of target sequence to the representative sequence (--min-seq-id 0.97 -c 0.8 --cov-mode 1). Hecatomb maintains the size of each cluster in the annotation table as well as the counts normalised to the total number of high-quality reads per individual sample (normalized as percent of the non-host reads). Clustering settings are also easily adjustable in the Hecatomb configuration file.

At the end of this process, non-redundant sequences have been removed and the remaining sequences are free from non-biological (reagent) contaminants and likely host-sequences. These high-quality sequences are then used for *de novo* metavirome assembly and read-based annotation.

### Metavirome assembly

A unique feature of Hecatomb is that it completes both individual read and assembly-based analysis. The first step of the metavirome assembly module is individual sample assemblies using MEGAHIT [53] (Figure 2). Long-reads are not amenable to using short-read assemblers and are therefore assembled using Canu [54]. Contigs from individual sample assemblies are subsequently assembled into a population assembly using Flye [64]. Per sample contig abundance are calculated by mapping individual sample reads to the population assembly. Read counts are reported normalised to library size and contig length using a variety of measures (reads per kilobase million (RPKM), fragments per kilobase million (FPKM) and sequences per million (SPM)). SPM is the same calculation as used for transcripts per kilobase million (TPM) except that the sequences are not assumed to be transcripts, thus the nomenclature adjustment. Calculations for RPKM, FPKM, and SPM are summarised in Supplementary Methods and an explanation is available in [65]. Taxonomic assignment of contigs in the population assembly is accomplished using MMseqs2 [41], queried against the secondary nucleotide database. Contig properties (e.g. length, GC-content) are combined with taxonomic assignments and sample abundance estimates into a final table. This contig table is merged with data obtained through the read based analysis to supplement contig mapping data with read-based taxonomic assignments and individual read properties.

**Figure 2:**
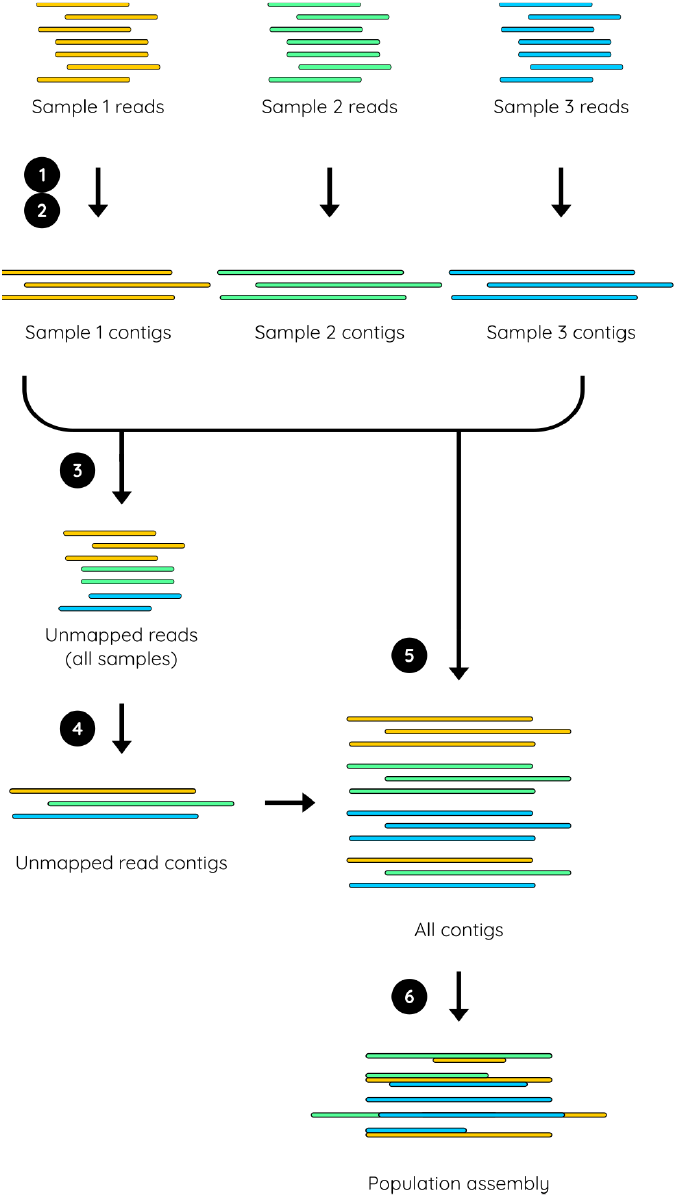
Metavirome Assembly. **(1)** High-quality kmer-normalised sequences from individual samples are assembled using either MEGAHIT or Canu. **(2)** The sequences for each sample are mapped to their respective assemblies. **(3)** The unmapped reads from all samples are pooled together. **(4)** The pooled unmapped reads are assembled using either MEGAHIT or Canu. **(5)** The contigs from all sample assemblies and the unmapped reads assembly are combined together. **(6)** Overlapping contigs are joined together using Flye using the subassemblies algorithm.

### Read-based annotation

Taxonomy (and functional information when available) is assigned using an iterative query strategy against both amino acid and nucleotide reference databases (Figure 3A). This strategy is designed to minimise false-positive viral annotations while maintaining sensitivity and runtime performance. All queries are carried out using MMseqs2 [41]. The strategy starts with a translated query of all sequences against a database of all viral (taxonomy ID: 10239) amino acid sequences in UniProtKB [66] clustered at 99% identity to reduce redundancy and target database size. Any sequence that matches a known viral protein is subsequently cross-checked against the complete multi-kingdom UniClust50 amino acid database [67]. The use of the well-annotated UniClust50 database enables functional as well as taxonomic annotation. This two-step query strategy captures all potential viral sequences in the first step, reducing the number of queries required in the secondary confirmatory step against the larger multi-kingdom database. The MMseqs2 searches can be time-consuming. Options are provided to use the default slower high-sensitivity parameters (--start-sens 1 --sens-steps 3 -s 7), or fast search parameters (-s 4.0) that yield greatly improved runtime performance.

**Figure 3:**
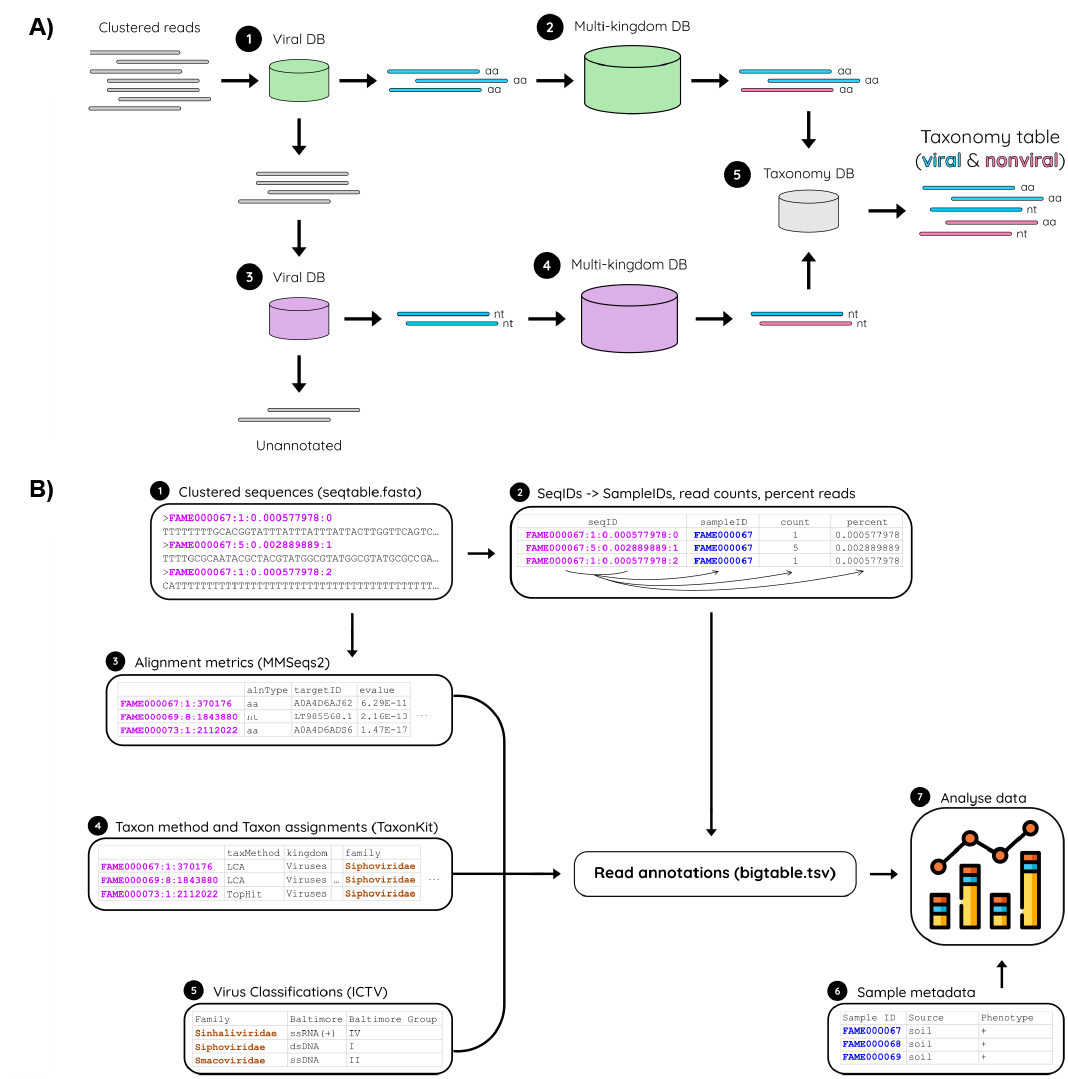
Read-based annotation. **a)** Iterative taxonomic annotation strategy. All alignments are completed using MMSeqs2. **(1)** High-quality representative sequences are queried against a viral amino acid (aa, *green*) sequence database. **(2)** Potentially viral sequences are subjected to a secondary, confirmatory query against a multi-kingdom amino acid sequence database. **(3)** Representative sequences that do not match a known viral amino acid are subjected to an untranslated query to a viral nucleic acid sequence database (nt, *purple*) **(4)** followed by a secondary, confirmatory query against a multi-kingdom nucleotide database **(5)**. Sequences that have been classified as either viral (*blue*) or nonviral (*pink*) in either the translated (aa database) or untranslated (nt database) queries are combined into a final taxonomy table. **(b)** Read annotation data structure. **(1)** Read Annotations are generated using the clustered sequences (seqtable.fasta). **(2)** The clustered sequence IDs are unpacked to yield the sample ID, the number of reads that sequence represents, and the percent of host-removed reads that sequence represents. **(3)** The alignment metrics from the annotation module are joined into the read annotations using the sequence ID as the primary key. **(4)** Taxonomic annotations are calculated and joined into the read annotations again using the sequence ID. **(5)** ICTV viral classifications are joined into the read annotations by the Taxonomic Family annotation. **(6)** Sample metadata can be joined into the read annotation table using the sample ID as the primary key. **(7)** The read annotation table with sample metadata can be quickly and easily analysed.

Sequences not identified as viral-like using translated queries to the amino acid database are subject to a similar iterative search using untranslated queries against a viral nucleotide sequence database consisting of all viral sequences in GenBank clustered at 100% identity to remove redundancy. This primary search is followed by a secondary confirmatory query against a polymicrobial nucleotide database containing representative RefSeq genomes from bacteria (n = 14,933), archaea (n = 511), fungi (n = 423), protozoa (n = 90) and plant (n = 145) genomes [68]. This iterative strategy enables sequence queries to target databases to be run on commodity hardware while still having representation of a broad diversity of non-viral kingdoms to minimise false-positive annotations.

Following secondary translated and untranslated searches Hecatomb augments sequence annotations using the lowest common ancestor (LCA) 2b-LCA algorithm described in [69]. This approach provides conservative taxonomic assignments at lower-nodes of the tree when similarity is found across a heterogeneous collection of taxonomies. However, the LCA algorithm fails when crossing higher taxonomic levels. For example, sequences with similarity to both bacterial and viral taxa have a LCA of “root” in the NCBI tree, while viruses from distinct viral domains are assigned to “virus root”. Hecatomb detects these instances and instead of classifying them to the root lineages refactors to the top-hit annotation. While this sometimes results in the reclassification of sequences to a non-virus lineage (e.g. if the tophit was bacterial) this novel approach provides additional information about sequences with ambiguous taxonomic assignments. This can be useful for instance in the identification of prophage regions which remains a challenging area of research [70,71].

### Outputs

Hecatomb output files are described in Supplementary Methods. Output tables are all tab-separated value (.tsv) files to ensure ease of use with data analysis. This tabular format is universally compatible with commonly used research software and programming languages such as Python, R, Excel or Bash and is easily merged with data from external sources, such as viral Baltimore classifications, International Committee on Taxonomy of Viruses (ICTV) taxonomy, or other external sources. The read annotation file is designed to acquire, preserve and organise data obtained throughout the pipeline with both study specific sample information as well as external data sources (Figure 3B). The process of investigating and removing false-positive annotations in viral metagenomes can be complex, but the abundance of alignment metrics in this file is designed to empower researchers to perform this step quickly and easily.

## Results

### Re-evaluation of a mammalian host-associated enteric virome

Hecatomb’s data structure (Figure 3B) integrates a large amount of information about individual sequences including taxonomic lineages, alignment statistics (e.g. E-values, percent identity, alignment length) and data from external virus information resources (e.g. Baltimore classification). To assess how this data structure can be used to evaluate the content of a complex virome we reanalysed a previously published data set (95 samples) obtained from stool samples collected from SIV-infected rhesus macaques (*Macaca mulatta*) (NCBI BioProject accession: PRJEB9503) [16]. Sequence data were generated using the Illumina MiSeq 2×250 bp paired-end protocol on libraries of total nucleic acid (DNA and cDNA to enable detection of both RNA and DNA viruses) extracted from stool samples. This data set was selected as it contains sequences from viruses from multiple Baltimore classifications (RNA and DNA genomes) that infect a variety of cell types (e.g. animal and plant). In addition, the original study identified differences in enteric virus abundances associated with SIV infection, enabling a comparative quantitative benchmark to evaluate Hecatomb with previously published results.

For the reevaluation study, Hecatomb was run using default parameters. Hecatomb’s taxonomic assignments classified sequences into phylogenetically diverse groups (Figure 4A). Bacteriophage from the family Microviridae and the order Caudovirales, (Siphoviridae, Myoviridae and Podoviridae), were the most abundantly classified viral sequence in the study. Hecatomb also identified a large number of sequences belonging to the Picornaviridae and Adenoviridiae, viral families regularly associated with gastrointestinal disease. Picronaviruses and adenoviruses were also identified in the original study with several adenoviruses having their full genomes sequenced as well as plaque purified [72]. Hecatomb also classified sequences belonging to a diverse set of viruses typically associated with infection of plants and protists (Figure S1).

**Figure 4:**
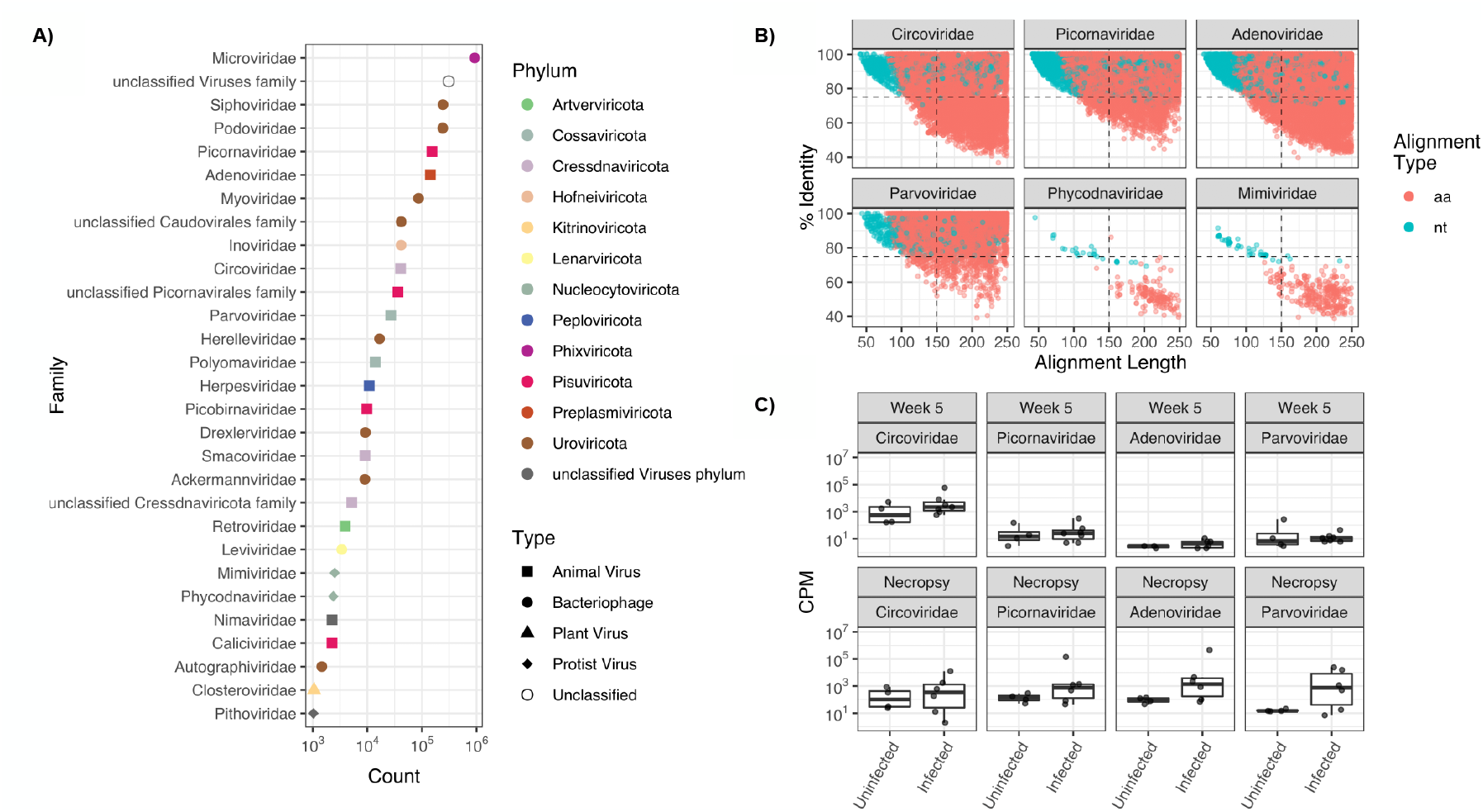
Reanalysis of rhesus macaque stool viromes. **(A)** Abundance of reads classified by viral Phylum (colour) and Type (shape). Phyla represented by fewer than 1,000 reads were excluded. **(B)** Percent identity and alignment lengths of all sequences classified for the 4 animal viruses identified in the previous study and two viruses of protists. Horizontal (70% identity) and vertical (150 base alignment length) dashed lines indicate a user-defined quadrant space. Each point represents an individual sequence colored by classification method (aa = classified via a translated search to an amino acid database, nt = classified via an untranslated search to a nucleotide database). Panels A and B represent data obtained from all 95 samples in the study. **(C)** Comparison of the number of sequences in SIV-infected and uninfected samples. Significance determined by the Wilcoxon signed-rank test. * = P ≤ 0.05, ** P ≤ 0.01, *** P ≤ 0.001, **** P ≤ 0.0001.

Hecatomb assigns NCBI taxonomy [73] using MMseqs2 [41] to query metagenomic sequences to relevant reference sequence databases. Taxonomic assignments relying on sequence similarity are dependent on the thresholds chosen. A permissive threshold risks increasing the rate of false-positives, while a stringent threshold may result in an increased rate of false-negatives. A perfectly accurate threshold is unlikely to exist, particularly given the high-variability in evolutionary histories across all viral types. In this case, plots and additional statistical analysis can prove useful in evaluating true- and false-positive viral annotations. Hecatomb collects alignment statistics (e.g. e-values, percent identity, alignment length, etc.) in the taxonomic assignment module and organises these data to assist in the identification of both true and false-positive taxonomic classifications.

As an example of how the alignment statistics can be used to evaluate true-or false-positive taxonomic assignments we examined percent identity and alignment lengths of the four viral families identified in the original study (Circoviridae, Picornaviridae, Adenoviridae and Parvoviridae). Hecatomb also annotated sequences to these same 4 viral families using both translated queries to amino acid (aa) databases and untranslated queries to nucleotide (nt) databases (Figure 4B).

While the statistics underlying sequence similarity searches are well understood, the application of thresholds to those statistics to infer taxonomy and function are more nebulous. Therefore, Hecatomb provides some additional guidelines to aid with the determination of true positives compared to false positives. For example, a quadrant system can be used to evaluate individual per family (or other taxonomic level) assignments (Figure 4B). Sequences in the upper two quadrants are highly similar to sequences in the reference databases over short (upper left, quartile 1 (Q1)) or long (upper right, Q2) alignment lengths, while sequences in the lower two quadrants have low similarity over short (lower left, Q3) or long (lower right, Q4) alignment lengths. For this analysis we arbitrarily selected 70% identity to represent the cut-off between low and high-identity for translated (aa database) and 90% identity for untranslated (nt database) alignments. Translated alignment length is reported in nucleotide base pairs rather than amino acid length. Therefore, a cutoff of 150 base pairs for both translated and untranslated alignment lengths was chosen (Figure 4B). Using this framework it is clear that there are many query sequences with high-identity (both short and long alignments) to sequences in both the aa and nt reference databases for the 4 families of previously identified animal viruses (Figure S2).

There were also a large number of query sequences classified as having statistically significant sequence similarity to reference sequences from viruses of protists (Figure 4B). Mimiviridae, that infect Acanthamoeba, and Phycodnaviridae, that infect algae, are both dsDNA viruses with large genomes [74]. While it is conceivable that these viruses may exist in the stool samples of rhesus macaques via water or food, using the quadrant framework there is little or no evidence of high-identity alignments to any sequence in either the aa or nt databases (Figure 4B, Figure S2). Hecatomb does not automatically remove sequences from these families as they would be common in environmental datasets. There is evidence for short and long low identity alignments (quadrant 4) to both Phycodnaviridae and Mimiviridae reference sequences. Thus, these sequences should be analysed using additional metrics (i.e. E-values, abundance across samples, etc.) to determine if these represent potentially novel viral sequences. This would not have been possible using stringent E-value filtering prior to data analysis.

Hecatomb also quantifies the normalised number of sequences (percent of host-removed reads) at each taxonomic depth. The normalised percent abundances per sample can be evaluated as the number of sequences assigned to a taxonomy per sample enabling statistical comparisons. The original study found evidence for four families of animal viruses (Circoviridae, Picornaviridae, Adenoviridae and Parvoviridae) in stool samples obtained from macaques infected with SIV or uninfected controls. The abundance of sequences from each viral family were similar between SIV-infected and uninfected macaques early in the study, but the abundance increased significantly as SIV-infection progressed while remaining the same in uninfected control animals. Evaluation of the normalised abundances for each of these four viral families using Hecatomb confirmed the findings of the original analysis (Figure 4C).

There were several viral families represented only using untranslated alignment to Hecatombs nucleotide database, including the Herpesviridae (Figure 5). All of the sequences assigned to the Herpesviridae aligned to only three target GenBank entries (Figure 5B). One entry (AF191073) dominated the similarities. All three were assigned a taxonomy with very low E-values suggesting statistically significant alignments (Figure 5C). However, all three of these entries belong to a single type of herpesvirus, Stealth virus 1 clone 3B43 [75]. The Stealth virus 1 genome was originally described as containing sections of both bacterial and viral genes. The three Stealth virus sequences identified by Hecatomb are identical to the bacterial segments when queried against the NCBI nt database (Figure 5D), suggesting that they are bacterial in origin. Very few sequences were found with alignments to the viral portion of the Stealth Virus 1 genome, which would be expected due to the random, shotgun sequencing process. This suggests that these sequences were called viral by hecatomb due to their similarity to a bacterial region of a viral genome, but that they are more likely bacterial false-positive contamination. Indeed, the original study identified Herpesviridae and many other false-positive sequences that were only removed following computationally-expensive blastn and blastx searches of the Non-Redundant nucleotide and protein databases [76].

**Figure 5:**
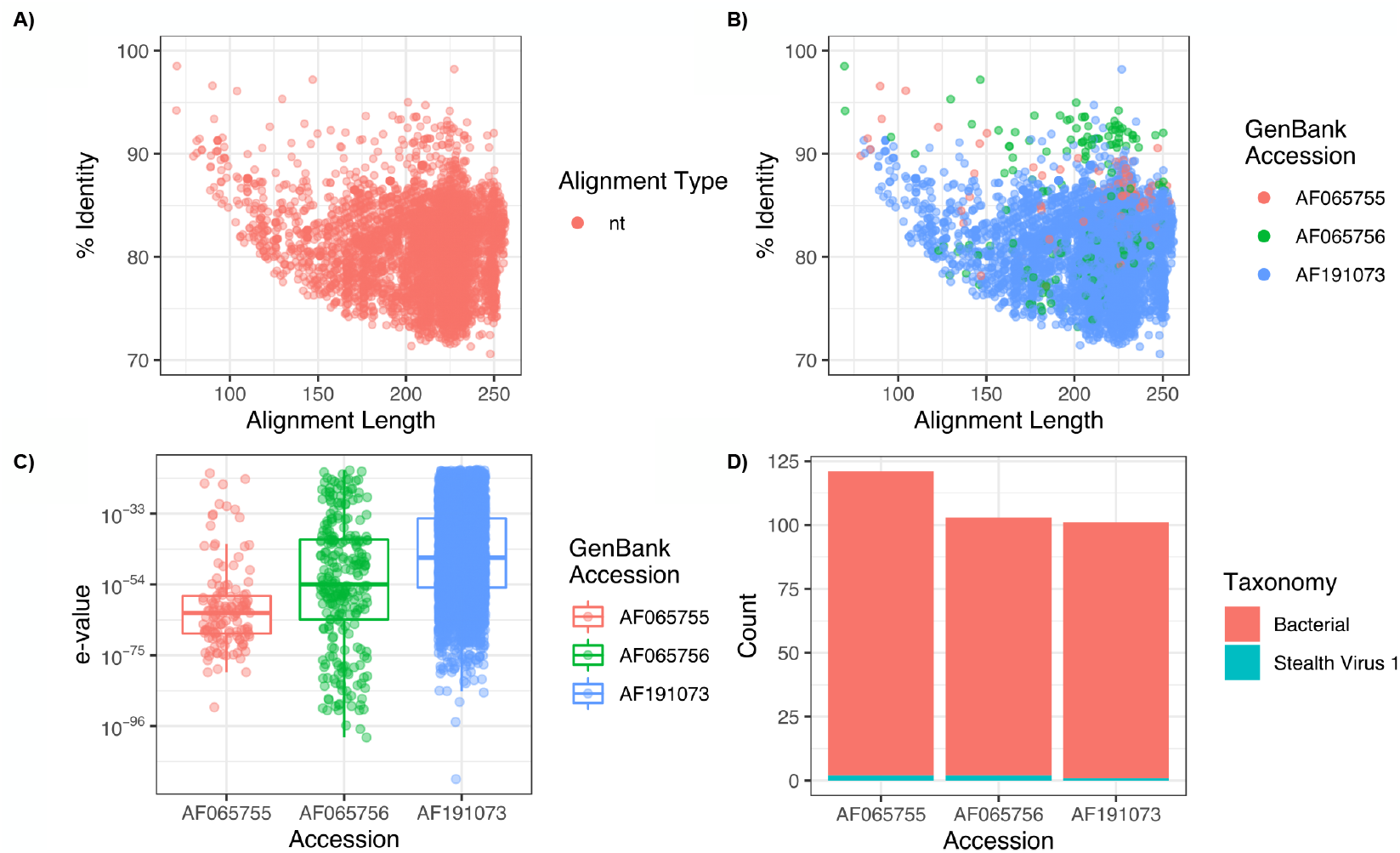
Ambiguous classification of bacterial sequences as Herpesviridae. **(A)** Percent identity and alignment length of all sequences assigned to the Herpesviridae.Note, there are no reads that were assigned using a translated search to an amino acid (aa) database. **(B)** Representation of GenBank accessions assigned to the Herpesviridae. **(C)** Summary of e-values for the 3 Herpesviridae accessions. **(D)** Summary counts of the taxonomic hits using blastn to the NCBI nucleotide (nt) database for each accession.

### Evaluation of an environmental dataset

We assessed Hecatomb’s ability to analyse non-human associated viromes by processing a previously studied coral reef dataset (NCBI BioProject accession: PRJNA595374) [77,78]. The dataset consists of metagenomic sequencing (Illumina MiSeq, paired 2×250) of both seawater and coral mucus from inner and outer sections of a Bermuda reef system. The original studies identified statistically significant differences in bacterial compositions between the coral mucus and seawater microbiomes and the coral mucus microbiomes from the inner and outer reefs. However, the viruses were not described in the original study. To further interrogate the viruses in these samples, study sequences were downloaded from SRA and run through Hecatomb using the fast parameters (--fast). Of the top 20 most abundant viral families, 10 are bacteriophages (Figure 6A). The relative abundance of viral families are mostly higher in inner reef samples compared to outer reef samples with exceptions such as Herelleviridae, Adintoviridae, Inoviridae, and unclassified Cressdnaviricota.

**Figure 6:**
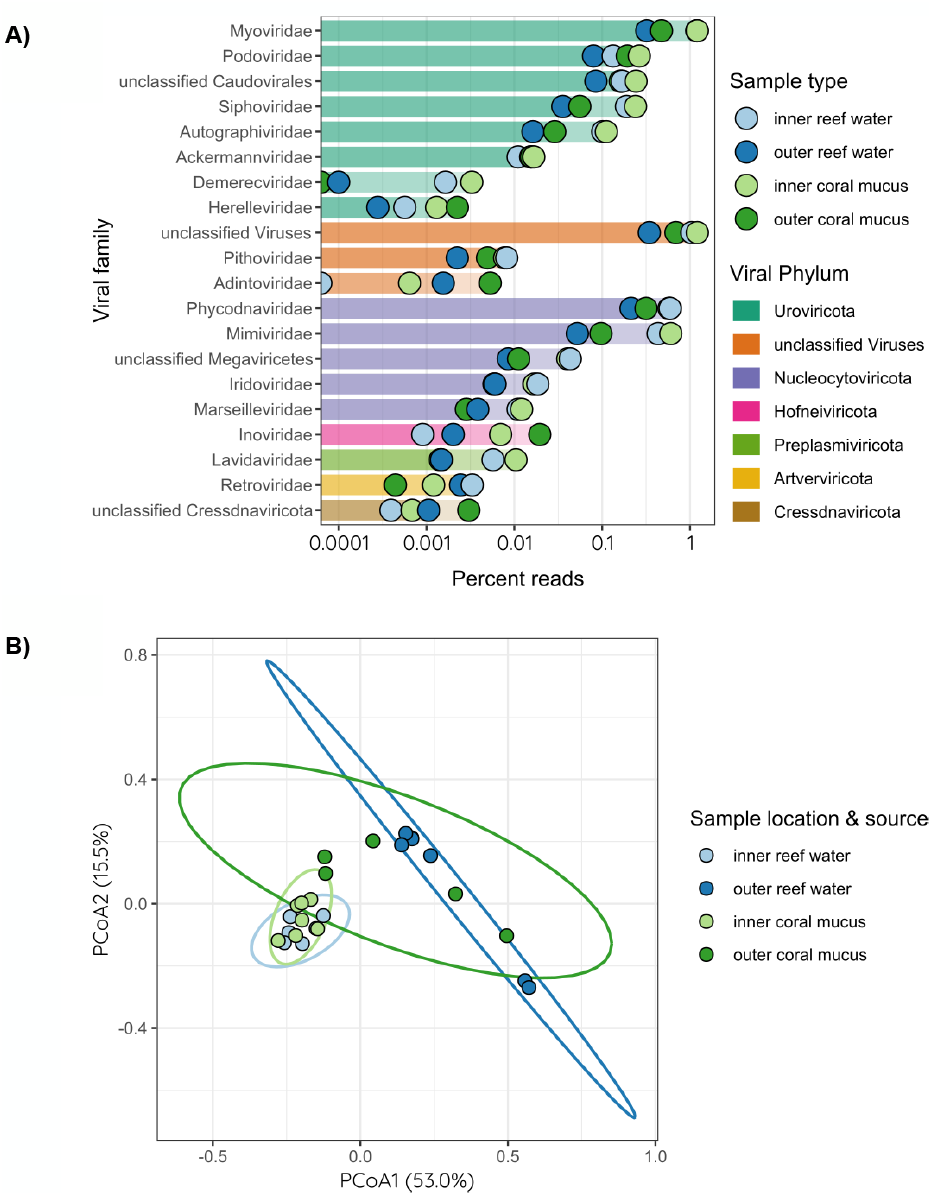
Reanalysis of Coral Reef Metagenomes. **(a)** The 20 most abundant viral families across coral reef samples. The sum of percent reads for each sample type are shown for each viral family. Viral families have been ordered and coloured by their Phyla. Points have been colored by sample type with inner and outer reef water samples coloured light- and dark-blue respectively, and inner and outer coral mucus samples colored light- and dark-green respectively. **(b)** Principle coordinate analysis (PCoA) of viral species abundance. Inner and outer reef water samples are coloured light blue and dark blue respectively. Inner and outer coral mucus samples are coloured light and dark green respectively. The Vegan package was used to calculate a bray-curtis distance matrix from the viral species counts, followed by multivariate dispersions with betadisp, and an Analysis of Variance (ANOVA) identified a non-homogenous distribution (P = 0.053). Ellipses for sample groups are drawn at 95% confidence levels for multivariate t-distribution.

Principal coordinate analysis (PCoA) of Bray-Curtis dissimilarity and a subsequent analysis of variance (ANOVA) confirmed a non-homogenous distribution across samples and groups (p = 0.053) (Figure 6B). Inner reef samples cluster closely together whereas outer reef samples appear to be far more varied. To examine compositional differences at each site, we performed permutational analysis of variance (PERMANOVA) of Bray-Curtis dissimilarity. We find that samples differ based on position (inner vs. outer reef: p = 0.001) and on the combined position and source (reef and mucus: p = 0.001), but not on source alone (inner vs. outer mucus: p = 0.185).

We calculated similarity percentage (SIMPER) between inner and outer samples, and between the outer reef samples only to identify viruses distinct to each group. SIMPER analysis identified many viral species that were significantly more abundant in inner reef samples, but none that were more abundant in the outer reef samples (Figure S3). In particular, Hecatomb classified reads to over 20 species of Synechococcus phage as being associated with outer reef samples. Viruses that contributed the largest fold differences included a phage that infects Verrucomicrobia (a mucin-degrading bacteria), and Namao virus (a Mimiviridae protozoan virus) which might infect Symbiodinium– coral’s endosymbiotic dinoflagellate.

When comparing outer reef coral mucus with outer reef water samples, we identified eight viruses that were more abundant in the reef water samples, with a phage that infects Halomonas bacteria as the largest fold difference (Figure S4). The largest fold differences observed in the coral samples included a Pyramimonas algae virus, a Vibrio phage, a Rhizobium phage, and a Pseudomonas phage.

## Discussion

Virome sequencing is the premier approach to evaluate the viral content of both host-derived and environmental samples. In the broadest terms, virome sequencing is used to answer two questions: i) What individual viruses are present in a sample or set of samples? ii) How does virome composition compare between groups of samples? The answers to these questions can be used to evaluate different biological questions. For example, knowing what individual viruses are present in a sample can be useful for identifying etiological agents of infectious disease. In contrast, analysis of the total virome or collection of viruses within a sample can be used to characterise ecological niches between groups. Both types of studies are dependent on effective computational tools not only to identify and classify viral reads within a metagenome, but also to assist in interpretation of complex virome data in association with study data.

Virome analysis is almost entirely dependent on sequence similarity queries against reference sequence databases. Historically, there have been two approaches to accomplishing this. The first is ‘brute force’ wherein all unclassified sequences are queried against a comprehensive, multi-kingdom reference sequence database (e.g. NCBI nt or nr). This approach relies on the search algorithm (e.g. BLAST, DIAMOND [79]) to pick the best or lowest-common ancestor of a group of hits to provide a final taxonomic assignment to an unknown query sequence. Hecatomb takes a different approach by first capturing all ‘potentially viral’ sequences by first querying sequences against a viral sequence database. These ‘potentially viral’ sequences typically represent only a small fraction of the full metagenomic data making subsequent computation more tractable. To confirm viral taxonomic assignment, all potentially viral sequences are cross-checked against a curated small transkingdom reference database containing genomic representatives from all kingdoms of life. Hecatomb completes this iterative search approach using translated searches against amino acid databases as well as untranslated searches against nucleotide databases, combining the results of each to ensure detection of viral sequences is database independent. This iterative search strategy uses databases orders of magnitude smaller than comprehensive, multi-kingdom databases (such as nt and nr) increasing computational efficiency without limiting viral detection.

Hecatombs’ design philosophy recognizes that there are no ‘perfect’ databases or search algorithms. Both the brute force and iterative search approaches against comprehensive or curated databases will result in different rates of true/false positives/negatives. Instead, Hecatomb relies on providing a compiled and rich set of data for search result evaluation. We used this strategy to reassess the virome composition of SIV-infected and uninfected rhesus macaques [16]. The original study used an iterative approach, but relied on comprehensive, transkingdom databases (NCBI nt and nr) and identified associations between four families of animal viruses (Circoviridae, Picornaviridae, Adenoviridae and Parvoviridae) and SIV-infection. The new Hecatomb trans-kingdom database is 6 orders of magnitude smaller than GenBank nt (5.0×10^6^ versus 1.3×10^12^) which results in a significant reduction in computational time and resources. Hecatomb identified the same four viral families and their relationship to SIV mediated disease (Figure 4C). Similar to our analysis of these samples using Hecatomb, the original study also classified a number of sequences to the Mimiviridae and Phycodnaviridae. Statistical comparison of these sequences between groups (e.g. SIV-infected vs. uninfected) did not reveal any significant associations thus they were not discussed further. However, new evaluation of results from Hecatomb indicates that there were likely false-positive classifications reported in the original analysis (Figure 4B). This highlights how coordinated data such as alignment statistics and taxonomy can be powerful tools for virome evaluation.

We were also able to evaluate the viromes of environmental (non-host associated) viromes. This analysis was primarily designed to identify compositional changes in viromes between reef types (inner or outer) and within coral mucosa and the surrounding water from a previously published metagenomic data set [77,78]. The original study identified elevated levels of Pelagibacter, Synechococcus, and unclassified Rickettsiales in inner reef samples compared to outer reef samples. Indeed, we found many elevated Synechococcus phages and other cyanophages in inner reef samples. However, we found only a few Mimiviridae viruses that were elevated which might be associated with Pelagibacter and unclassified Rickettsiales, despite Pelagibacter being identified as the most abundant genus in the original study. It’s possible that Synechococcus and other cyanobacteria growth rates are high, and that this is offset by greater viral activity (a viral shunt) that results in nutrient cycling to other microbes in the reef system. Heterotrophic bacteria and archaea are significant sources of fixed-nitrogen in coral reefs (reviewed in [80]), so viral activity of cyanobacteria would therefore be beneficial to the entire reef ecosystem by supplying both organic nitrogen, and by feeding these nitrogen-fixing bacteria.

The inner reef coral mucus and reef water viromes clustered tightly suggesting that there was little difference in these viromes. The consistency in viral compositions between coral mucus and reef water samples of the inner reef systems is interesting and suggests an equilibrated flux of viral particles between coral mucus microbiomes and the surrounding reef water. Conversely, differences were observed in viral abundances of outer reef samples, and most were found to be species that were more elevated in coral mucus compared to reef water samples. The greater differences in viral compositions between the outer reef coral mucus and water samples could indicate that the greater exchange of water between the reef system and open ocean may be depleting viruses from this ecosystem. Furthermore, the greater thermal stability and reduced particulate load (from terrestrial runoff) results in a reduced turnover of coral mucus in the outer reef samples (described in [77,78]), which may also contribute to the higher relative abundances of viruses in inner reef systems in general.

## Conclusions

Virome analysis is complex and requires efficient computational tools to generate analyst friendly results. Hecatomb provides a comprehensive and computationally efficient solution for both read- and assembly-based viral annotation and virome analysis. The pipeline is delivered with a convenient and easy-to-use front end and is compatible with different sequencing technologies. Hecatomb’s comprehensive collection of data throughout the running of the pipeline, in particular the collection of alignment statistics, empowers identification and interrogation of viral taxonomic assignments. We demonstrate Hecatomb’s utility for rapid processing and analysis of viral metagenomes with a well-studied validation gut viral metagenome dataset. We also demonstrate its utility for mining regular metagenome samples for virome analysis by analysing an existing environmental dataset.

## Declarations

### Ethics approval and consent to participate

Not applicable.

### Consent for publication

All authors have confirmed consent for publication.

### Availability of data and materials

**Project name**: Hecatomb

**Project home page**: github.com/shandley/hecatomb

**Project documentation:** hecatomb.readthedocs.io

**Operating system**: Linux

**Programming language**: Python

**Other requirements**: Conda

**Licence**: MIT

### Restrictions to use by non-academics

None

The reanalysis with Hecatomb utilised pre-existing datasets which are available under the NCBI BioProject accessions PRJEB9503 for the macaque SIV dataset [16] and PRJNA595374 for the coral reef dataset [77,78]. The Hecatomb annotations are available at doi.org/10.5281/zenodo.6388251, and all commands used for analysing the results are available at gist.github.com/beardymcjohnface/3d3245b2bf6d9544c524f412037d5065.

### Author’s contributions

MJR, RAE, and SAH conceived the pipeline and data structures. KAM, LW, and AP provided suggestions about the pipeline and data visualisations. MJR, SJB, RAE, and SAH coded the pipeline. KH-C contributed to documentation and analysis. MJR and SAH performed the analysis and interpretation. LFOL, RAE, and EAD helped with interpretation of results. MJR, RAE, and SAH drafted the original manuscript. All authors reviewed and edited the manuscript.

## Acknowledgments

The authors thank Chandni Desai and Barry Hykes for their thoughtful commentary regarding the design philosophy of Hecatomb, and Sarah Giles, Susie Grigson, Bhavya Papudeshi, Vijini Mallawaarachchi, and Laura Inglis for feedback on the manuscript. The support provided by Flinders University for HPC research resources is acknowledged.

## List of abbreviations

AIDS: acquired immunodeficiency syndrome
SIV: simian immunodeficiency virus
HPC: high-performance computing
NCBI: National Center for Biotechnology Information
RPKM: reads per kilobase million
FPKM: fragments per kilobase million
SPM: sequences per million
LCA: lowest common ancestor
ICTV: International Committee on Taxonomy of Viruses
PERMANOVA: permutational analysis of variance
PCoA: principal coordinate analysis
ANOVA: analysis of variance
SIMPER: similarity percentag

## Figure Legends

**Figure S1:Taxonomic subsets of virus types**

Viral families present in the 95-sample SIV reanalysis study **(A)** Plant viruses, and **(C)** Protist viruses

**Figure S2:Sequence per Quadrant Evaluation**

Percentage of reads per quadrant in Figure 5. **(A)** translated (aa reference database) and **(B)** untranslated (nt reference database)

**Figure S3:Viral abundance for inner and outer reef samples**

Viruses more abundant by Similarity Percentage (SIMPER) analysis in inner reef samples are colored red. Viral species constituting 95% of variance that are significantly different (p<0.05, log2 fold difference > 2) are shown. Infinite values are capped at an absolute log2 fold difference of 5.

**Figure S4:Viral abundance for outer reef coral mucus and outer reef water samples**

Viruses more abundant by Similarity Percentage (SIMPER) analysis in outer reef coral mucus samples and outer reef water samples are coloured red and blue respectively. Viral species constituting 95% of variance that are significantly different (p<0.05, log2 fold difference > 1) are shown. Infinite values are capped at an absolute log2 fold difference of 5.

