## Supplemental Material for "Hecatomb: An End-to-End Research Platform for Viral Metagenomics"

### Supplementary Methods

**Hecatomb databases.** Hecatomb databases are hosted in the cloud with Amazon Web Services (AWS). Downloading and installing them is managed by the Hecatomb launcher. The most up to date descriptions of the databases can be obtained from Hecatomb's documentation at hecatomb.readthedocs.io.

1. **Contaminants**. This database consists of a collection of NEBNext and TruSeq sequencing adapters, primers, and vector contaminants from UniVec [62]. The contaminants database is used exclusively during sequence preprocessing.

2. **Hosts**. A collection of host genomes that have been preprocessed to mask viral-like and low-entropy sequences. Host genomes are used for host read removal during preprocessing. Hecatomb comes with several common host genomes ready to use, and users can add their own host genomes to the database via the Hecatomb launcher.

3. **AA and NT**. These are the amino acid (AA) and nucleotide (NT) databases used for sequence annotation of both reads and contigs. For each of the AA and NT databases, there is a primary viral database used for classifying reads that match a known virus, and a secondary multi-kingdom database which is used for assigning taxonomy to either reads and contigs. The primary AA database includes all UniProt viral protein entries clustered at 99% identity. The secondary AA database consists of the Uniclust50 database [67] (doi.org/10.1093/nar/gkw1081) supplemented with the primary AA database. The primary NT database consists of all viral sequences in GenBank clustered at 100% identity to remove

redundancy. The secondary NT database consists of a customised polymicrobial nucleotide database containing representative RefSeq genomes from Bacteria (n = 14,933), Archaea (n = 511), Fungi (n = 423), Protozoa (n = 90) and plant (n = 145) genomes.

4. **Tax**. This is NCBI's taxonomy database used for translating taxonomy IDs into full taxonomic lineages [73].

5. **Tables**. These are additional classification tables such as the ICTV 2019 Baltimore groups for viruses, which are used to supplement the output annotation tables.

**Calculations for RPKM, FPKM, and SPM**

*Reads per kilobase million (RPKM) calculation:*

$$(C \div (T \div 1{,}000{,}000)) \div L$$

$C$ = number of reads mapped to a contig

$T$ = total number of reads in a sample

$L$ = length of contig in kilobases

*Fragments per kilobase million (FPKM) calculation:*

Same as for RPKM except that paired reads are only counted as a single mapped read.

Sequences per Million (*SPM) calculation:*

$$(Ci \div Li) \div (\sum_{n} (Cn \div Ln) \div 1{,}000{,}000)$$

$Ci$ = number of reads mapped to a contig

$Li$ = length of a contig in kilobases

$\sum_{n} (Cn \div Ln)$ = sum of number of reads mapped to contigs divided by contig length

**Hecatomb outputs.** The most up-to-date descriptions of all the pipeline output files can be obtained from Hecatomb's documentation at hecatomb.readthedocs.io. A tutorial is included to demonstrate typical analyses based on these output files using both R or Python. The main output files are:

1. **Report**. Hecatomb utilises a Snakemake feature to generate a report (*report.html*) of the pipeline run. The report includes the pipeline rule graph, details about resource usage and runtime for each rule, and the used configuration file. The report also includes read counts generated following every preprocessing and annotation step rendered as a Sankey diagram.

2. **Seqtable**. The sequence table (*seqtable.fasta*) contains all representative sequences for the clustered reads for all samples. The sample names and cluster

counts are incorporated into the sequence IDs whilst remaining in a standard sequential multi-fasta format.

3. **Read annotations**. The read annotations table (*bigtable.tsv*) is used for the majority of analyses from Hecatomb. This table combines the sequence and sample IDs, cluster counts, alignment metrics, taxonomic and available functional annotations into one table. This table is designed to expedite downstream analysis of the read annotations using commonly used software platforms (Python, R, BASH, Excel).

4. **Assembly**. The population assembly (*assembly.fasta*) is the output of contigs obtained by Flye. Contigs are annotated using MMseqs2 queries against viral and other genomes obtained from the secondary nucleotide database [68].

5. **Contig annotations**. Contig taxonomic annotations (*contigAnnotations.tsv*) are calculated using MMSeqs2 [41] querying each contig against Hecatombs' multi-kingdom NT database (described above). Read-based contig annotations (*contigSeqTable.tsv*) are also generated by mapping individual read annotations positionally to each contig.

6. **Contig coverage**. Coverage statistics of each population assembled contig (*contig_count_table.tsv*) are calculated by mapping reads from each sample to all contigs using BBMap (https://sourceforge.net/projects/bbmap/). RPKM, FPKM, and SPM are subsequently calculated (see above Supplemental Methods for calculations). A read-based contig annotations file (*contigSeqTable.tsv*) can also be used to retrieve mapped read counts of contigs for each sample.

**Table S1. Software packages used by Hecatomb.**

| Package name | Version | URL | Reference |
|---|---|---|---|
| Conda | 4.10.3 | conda.io | |
| Mamba | 0.17.0 | github.com/mamba-org/mamba | |
| Python3 | 3.10.0 | python.org | |
| Snaketool | 9d81e2b | github.com/beardymcjohnface/Snaketool | [49] |
| Snakemake | 6.4.1 | snakemake.github.io | [50] |
| fastp | 0.23.2 | github.com/OpenGene/fastp | [51] |
| PySam | 0.17.0 | github.com/pysam-developers/pysam | |
| BBTools | 38.90 | jgi.doe.gov/data-and-tools/bbtools | [52] |
| MEGAHIT | 1.2.9 | github.com/voutcn/megahit | [53] |
| Canu | 2.2 | github.com/marbl/canu | [54] |
| Flye | 2.7.1 | github.com/fenderglass/Flye | [55] |
| Minimap2 | 2.20 | github.com/lh3/minimap2 | [56] |
| Samtools | 1.13 | github.com/samtools/samtools | [57] |
| MMseqs2 | 12.113e3 | github.com/soedinglab/MMseqs2 | [41] |
| TaxonKit | 0.8.0 | github.com/shenwei356/taxonkit | [58] |
| Plotly | 5.3.1 | plotly.com | |
| Kaleido | 0.2.1 | pypi.org/project/kaleido | |
| SeqKit | 0.16.1 | github.com/shenwei356/seqkit | [59] |
