## Supplementary figures and images for "Hecatomb: An End-to-End Research Platform for Viral Metagenomics"

### Supplemental Figures

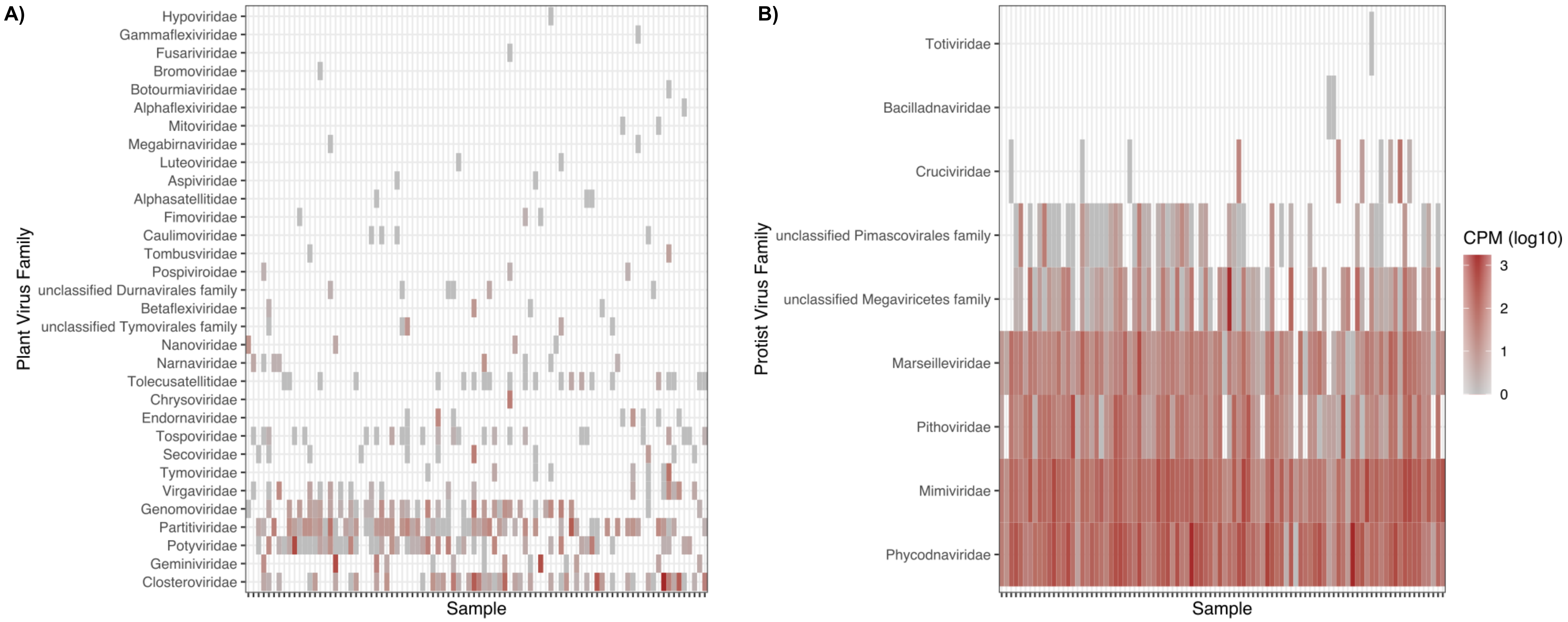

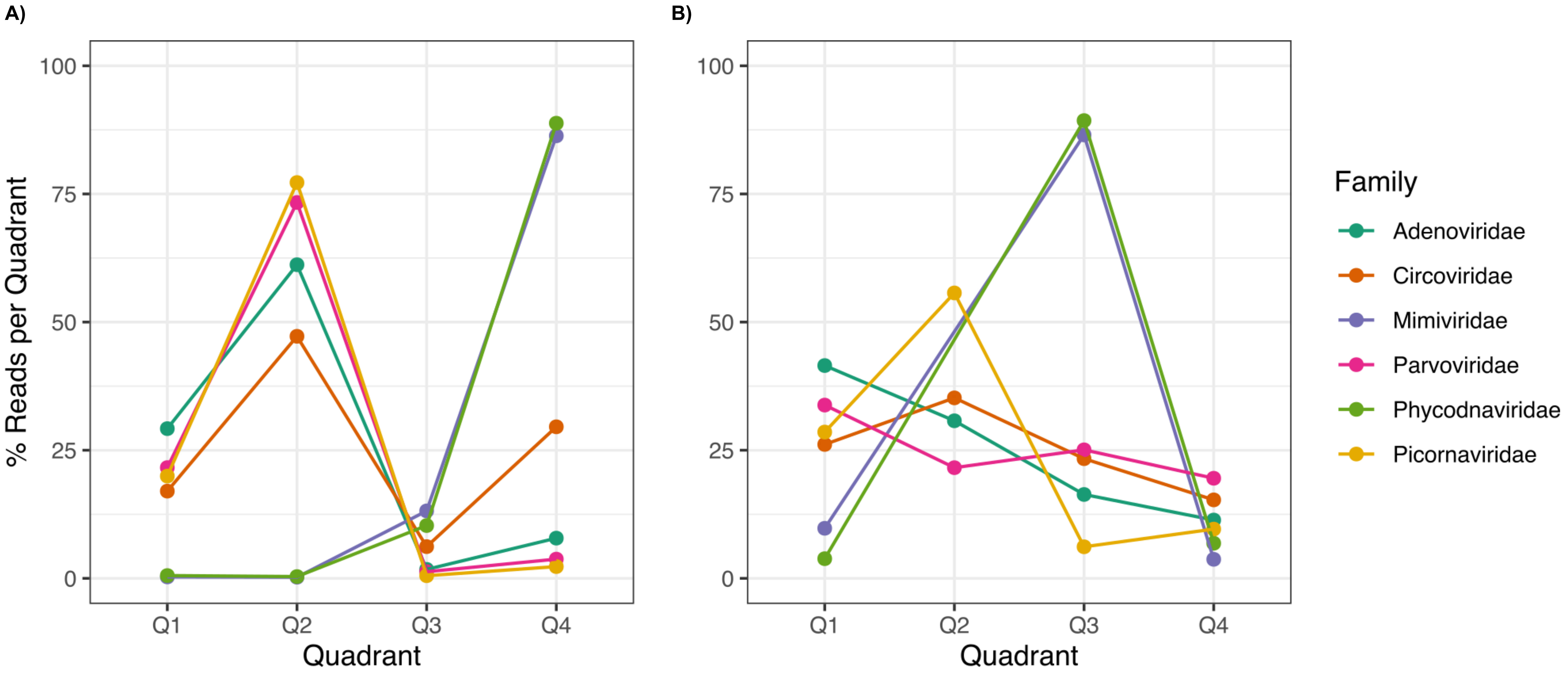

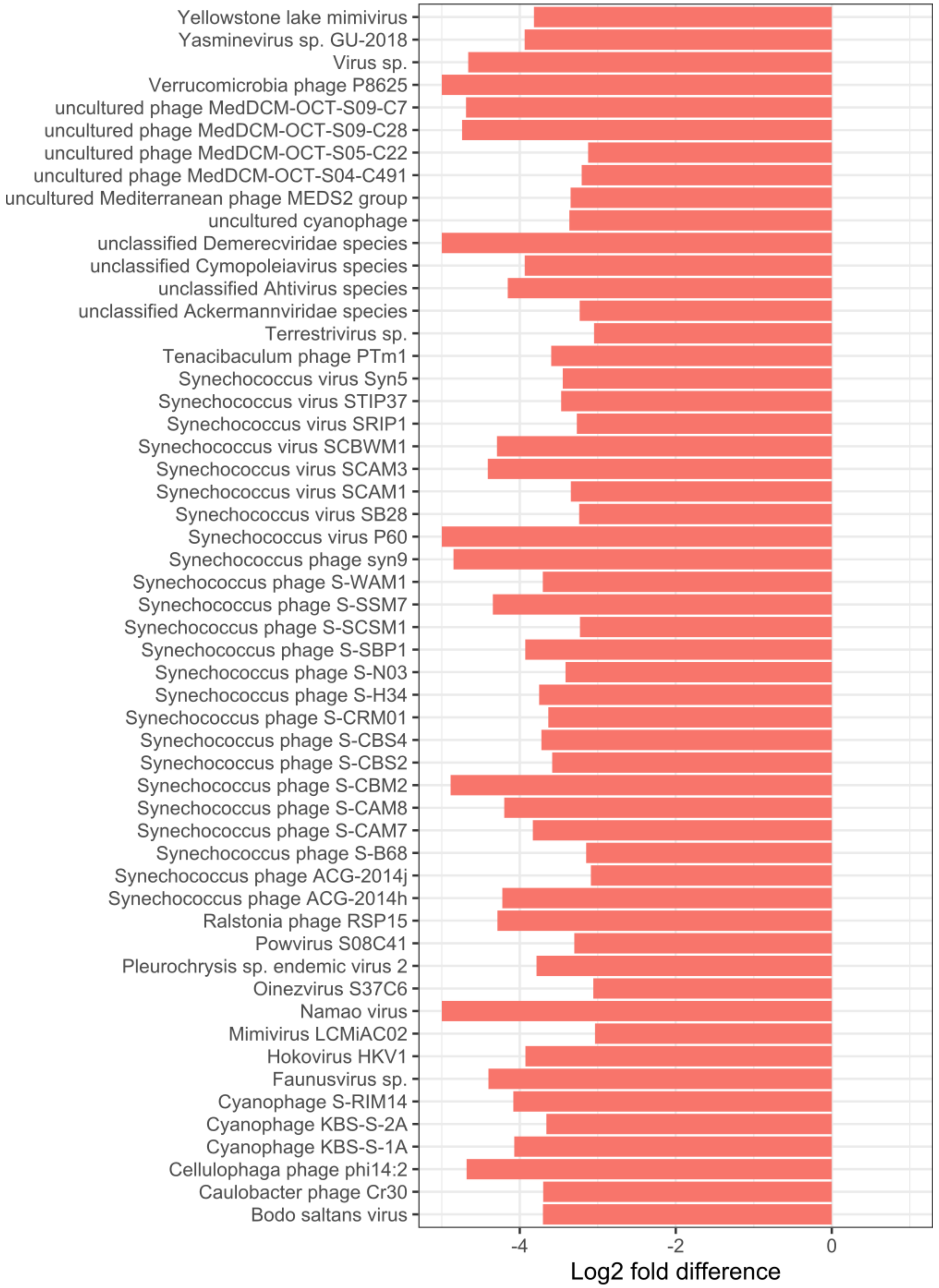

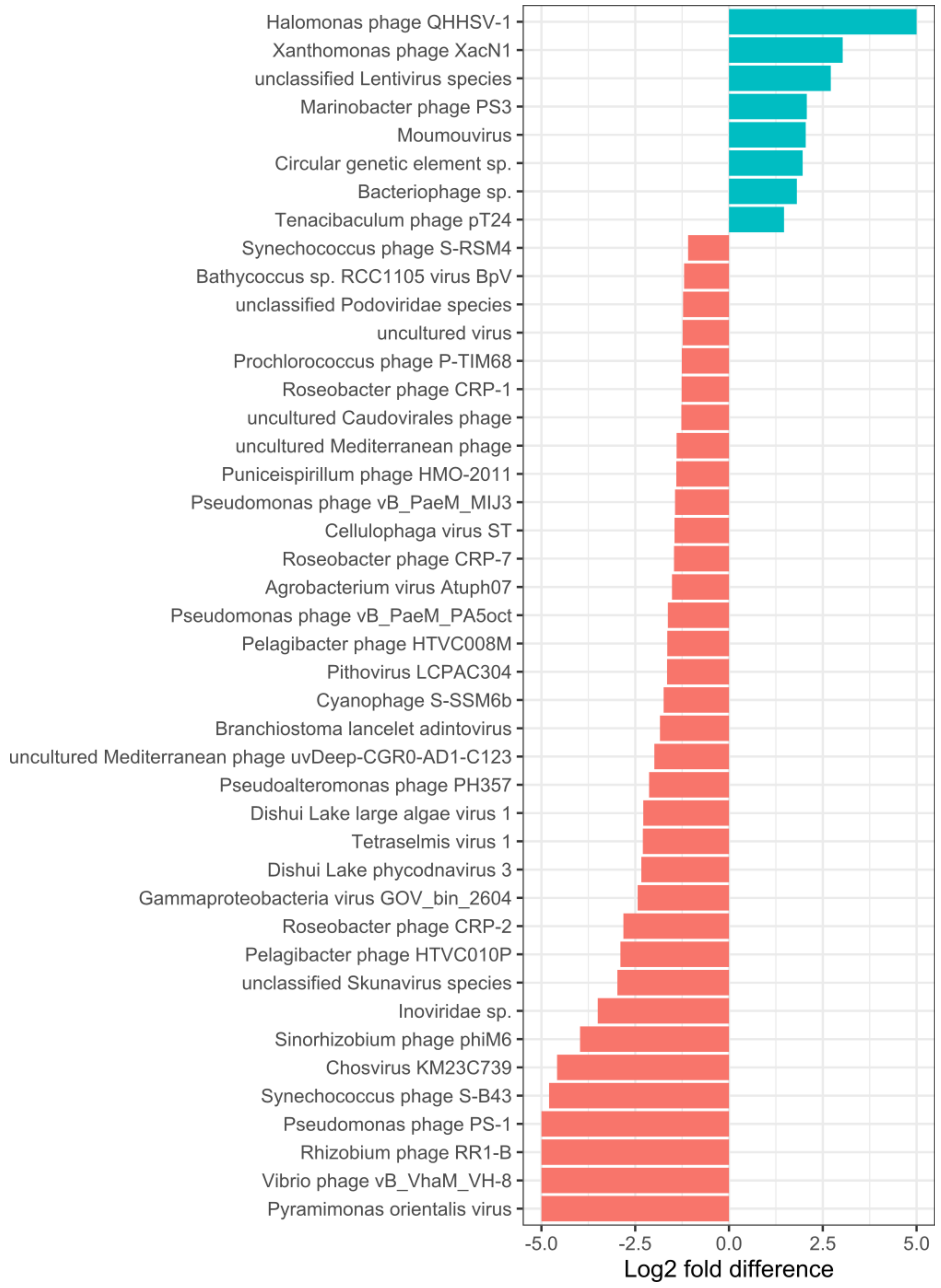
